# Shaft versus spine localization affects the structural plasticity of glutamatergic synapses

**DOI:** 10.64898/2025.12.05.692591

**Authors:** Tomas Fanutza, Yannes Popp, Arie Maeve Brueckner, Nathalie Hertrich, Judith von Sivers, Matthew Larkum, Sarah A. Shoichet, Marina Mikhaylova

## Abstract

In the early stages of development, most excitatory synapses are formed directly on dendritic shafts. As neurons mature, these sites gradually shift from the shaft to dendritic spines. In fully developed excitatory neurons, the majority of glutamatergic postsynaptic sites containing the postsynaptic density (PSD) molecules reside on dendritic spines. However, some glutamatergic synapses remain as shaft synapses, yet their characteristics have remained unexplored. Here, we show that the molecular composition of the shaft PSDs closely resembles that of spine PSDs. Key components such as AMPARs, NMDARs, Ca_V_1.2 channels, and F-actin interacting proteins, SynGAP, as well as cortactin, are present in comparable amounts in both synapse types. The major distinction between shaft and spine PSDs lies in the lower abundance of the scaffold proteins Shanks and Homer in shaft PSDs. Shaft synapses are not merely passive structures but actively participate in synaptic transmission. Their structure and function are modulated by changes in neuronal activity. Long-term live imaging combined with a cLTP protocol revealed that shaft PSDs were potentiated but rarely underwent a transition to spine synapses. In contrast, during LTD, shaft PSDs were eliminated more frequently than their spine counterparts. Together, these findings highlight excitatory shaft synapses as a distinct, and notably less stable, synapse type.

## Introduction

Neurons are highly specialized and compartmentalized cells that extend multiple dendrites that receive inputs and usually a single axon, which forwards information to the next cells.^1^ Neurons convey information via specialized contacts called synapses. One of the major classes of neurons in the central nervous system are excitatory glutamatergic neurons. They are highly abundant in the forebrain and integrate input from inhibitory and excitatory neurons across different brain regions. Inhibitory synapses are mostly formed on the dendritic shafts^2^, while in mature neurons, the majority of excitatory synapses are formed on specialized dendritic protrusions called dendritic spines.^3^ During early development, excitatory postsynaptic sites are almost exclusively located on the dendritic shaft. With maturation of the cell, the ratio shifts to the location on dendritic spines.^4^ However, 10-30 % of excitatory postsynaptic sites remain on the dendritic shaft.^3,5^

These excitatory shaft synapses have been described in a few studies using a variety of imaging techniques.^5–7^ As it was shown in primary hippocampal cultures, glutamatergic shaft synapses associate with a dense F-actin meshwork, called F-actin patches. This mesh consists of a mixture of longitudinal and branched actin filaments and, similarly to the F-actin in dendritic spines, surrounds the PSD of glutamatergic shaft synapses.^8^ Initial work has demonstrated that F-actin patches were adjacent to active presynaptic sites labeled with Bassoon, and synaptotagmin 1 (syt-1) antibody uptake assay.^8,9^ One of the identified functions of F-actin patches is stalling and positioning of endolysosomes next to the shaft synapses via the myosin V actin motor protein. In an *in vivo* study, all synapses on layer2/3 cortical pyramidal neurons were mapped using *in utero* electroporation and sparse labeling. Quantification of Homer1c-tdTomato puncta in brain sections of postnatal day 42 mice revealed that ∼10 % of excitatory postsynaptic sites were located on the dendritic shafts.^10^

A fraction of these sites can be accounted for by so-called silent synapses, which can be located on the dendritic shaft.^11,12^ Silent synapses are characterized by a PSD, which contains only N-methyl-D-aspartate (NMDA) receptors and lacks functional 2-amino-3-(3-hydroxy-5-methyl-isoxazol-4-yl) propanoic acid (AMPA) receptors. Super-resolution stochastic optical reconstruction microscopy (STORM) has unveiled that the silent synapses can be found on both spines and dendritic shafts.^6^

The main scaffolding components of the PSD are scaffold proteins like Homer, Shanks, and PSD95. They can bind to cytoskeletal elements and form nanodomains.^13,14^ They anchor synaptic membrane proteins like AMPA receptors, NMDA receptors, kainate receptors, and metabotropic glutamate receptors, as well as calcium channels and cell adhesion molecules (e.g., NCAM1, integrins, neuroligin-1).^15^ Importantly, scaffold proteins interact with the F-actin cytoskeleton via F-actin binding proteins α-actinin and cortactin. Therefore, changes in the F-actin network can influence the location and formation of postsynaptic sites and subsequently the expression of glutamate receptors and cell adhesion molecules. ^16,17^

Already more than two decades ago, it was shown that dendritic spines are not rigid structures and undergo rapid and constant remodeling that is dependent on F-actin dynamics.^18^ Spines can change their size and shape within seconds, enabling the neuron to rapidly adapt to changing demands for the postsynapse. This activity-dependent remodeling, referred to as structural plasticity, is a key mechanism through which synapses adjust their strength. Apart from the structural changes, the composition of the synapses can also change, modulating the synaptic transmission. Two of the most studied types of synaptic plasticity are long-term potentiation (LTP) and long-term depression (LTD).^19–21^ LTP results in a strengthening of the synaptic contacts. On the postsynaptic site, this can include an increase in the number of glutamate receptors, changes in the receptor subunit composition, and an increase in the size of PSDs. Contrary, LTD results in the removal of glutamate receptors from the PSD or the shrinkage and even disappearance of the PSD.^22,23^

Synaptic plasticity is required for complex processes like learning and memory. A study on learning in mice showed that within 4 days of learning a reaching task, the remodeling of spines in layer 5 cortical pyramidal neurons increased significantly.^24^ It was observed that both new spine formation and spine removal occurred more often compared to the control. Furthermore, the authors could show that newly created spines are sites of functional excitatory synapses.

While spine PSDs are well studied and characterized regarding their composition, plasticity, and function, the excitatory shaft postsynapses (shaft PSDs) are less investigated, and key questions remain open: Are these functional synapses? If so, do they undergo synaptic plasticity?

Here, we use organotypic rat hippocampal slice cultures and primary rat hippocampal neurons to investigate the abundance of excitatory shaft synapses in mature neurons, as well as their composition and activity in synaptic transmission, and compare them with spine synapses. Furthermore, we investigated whether shaft synapses can undergo structural plasticity upon induction of chemical LTP (cLTP) and chemical LTD (cLTD).

## Results

### Dendritic localization of glutamatergic postsynapses in CA1 hippocampal neurons

Hippocampal glutamatergic synapses in mature neurons exhibit a high degree of heterogeneity in terms of their molecular composition, structural features, and functional properties.^5,8,25^ To estimate the frequency distribution of excitatory shaft synapses in analogy to spine synapses, we employed single-cell electroporation to co-transfect mature CA1 pyramidal neurons with mRuby3 (cell fill) and PSD95.FingR-GFP (intrabody against endogenous PSD95)^26^ in organotypic rat hippocampal slices.

This allowed us to visualize and quantify the endogenous PSD95, which is a major scaffold protein, present in virtually all glutamatergic synapses (**Fig. 1**).^26–29^ At days *in vitro* (DIV) 21, organotypic slices were electroporated and kept for another 2 to 4 days, then fixed and imaged using high-resolution confocal microscopy. With this approach, expression of mRuby3 enabled clear identification of dendritic arborizations and distinction between two different morphological neuronal cell types with two different apical dendrite structures, namely bifurcated and unidirectional neurons (**Fig. 1A**). These two different cell architectures were previously reported by Benavides-Piccione and colleagues in neurons from both humans and mice.^30^

**Figure 1:**
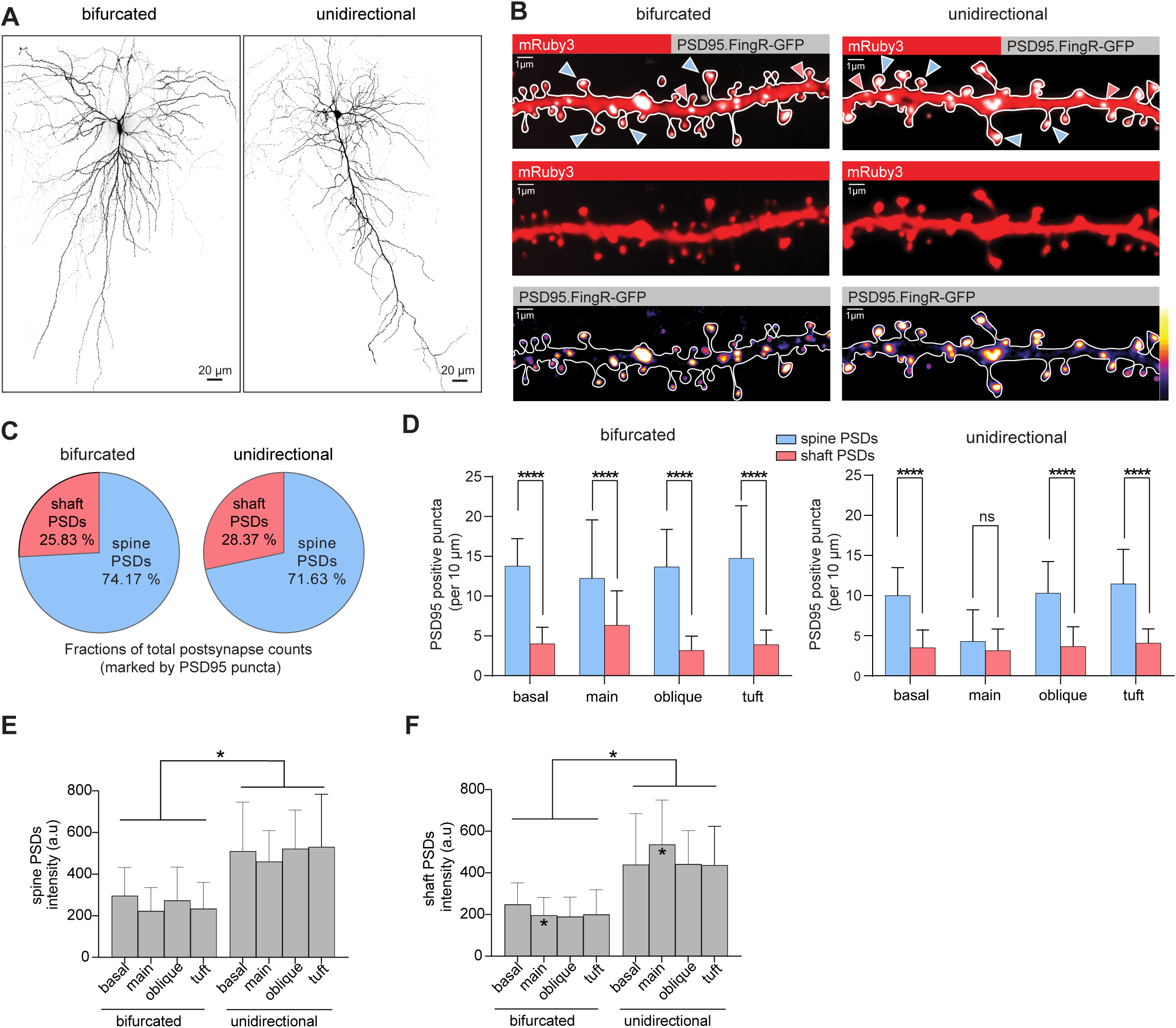
Glutamatergic synapses are predominantly located on dendritic spines, while some are located on the shaft of mature CA1 pyramidal cell dendrites. **A)** Representative maximum z-projections of confocal images showing apical dendritic structures of DIV21 CA1 pyramidal neurons, expressing mRuby3 (cell fill), with a branching (bifurcated) main shaft (left) and without branching (unidirectional) main shaft (right). **B)**Representative maximum z-projections of confocal images displaying dendritic segments from a bifurcated (left) and unidirectional (right) CA1 pyramidal neuron expressing mRuby3 (cell fill) and labeled for endogenous PSD95 (excitatory postsynaptic marker) by using the PSD95.FingR-GFP probe. Spine PSDs (blue arrows) and shaft PSDs (pink arrows) are indicated. Lower panel: pseudo color scale to visualize signal intensity differences of endogenously expressed PSD95. **C)** Quantification of spine and shaft PSDs in percentage of the total PSD95 puncta in bifurcated and unidirectional CA1 neurons. **D)** Quantification of spine and shaft PSDs by dendritic compartment in bifurcated (left) and unidirectional (right) neurons. Two-way ANOVA with Šídák’s multiple comparisons test. **** p<0.0001 (basal, main, oblique, tuft dendrites); p=0.4923 (main unidirectional dendrite). Data are presented as mean±SD. **E)** Quantification of intensity signals of endogenous PSD95 puncta in spine PSDs. Two-way ANOVA with Šídák’s multiple comparisons test; p=0.4331 (basal); p=0.0901 (main); p=0.2571 (oblique); p=0.1058 (tuft). Data are presented as mean±SD. **F)** Quantification of intensity signals of endogenous PSD95 puncta in shaft PSDs. Two-way ANOVA with Tukey’s multiple comparisons test; p=0.5127 (basal); p=0.0286 (main); p=0.1950 (oblique); p=0.2573 (tuft). Data are presented as mean±SD. All the quantifications have been performed on bifurcated neurons, from 27 dendritic segments in 8 cells. Unidirectional neurons, from 33 dendritic segments in 11 cells.

First, we quantified the number of dendritic protrusions in both neuronal cell types and calculated their densities. We found that dendrites of bifurcated neurons contain a significantly higher number of dendritic protrusions in comparison to unidirectional neurons (**Fig. S1B**). Second, the total number of excitatory postsynapses (PSD95-positive) on dendritic shafts (shaft PSDs) and spines (spine PSDs) across all dendritic compartments of bifurcated and unidirectional neurons was quantified (**Fig. S1A+C**). Consistent with the literature, PSD95 was expressed on most dendritic spines in both neuronal cell types (**Fig. S1F**).^29,31^ In addition, of the total number of PSD95 puncta counted, approximately 71-74 % were spine PSDs and 26-29 % were shaft PSDs (**Fig. 1B+C**). Despite these numbers suggesting there is a similar synaptic distribution, the quantification revealed overall a significantly higher number of PSD95-positive spines on bifurcated cells compared to unidirectional (**Fig. S1C**).

When observed by dendritic compartment, spine PSDs were significantly denser compared to shaft PSDs in both neuronal subtypes (**Fig. 1D**). The only exception was the main dendritic shaft of unidirectional neurons, where PSD95 puncta were evenly distributed between shaft and spine sites (**Fig. 1D**, right). Indeed, compared to bifurcated cells, the main dendritic shaft of unidirectional cells showed the most significant lack of spine PSDs, contributing largely to the overall lower spine PSD number compared to its bifurcated counterpart (**Fig. S1D+E**).

Our results, consistent with previous studies, revealed that excitatory synapses are predominantly formed on dendritic spines across all dendritic compartments, and we further demonstrated that cells with a bifurcated apical dendritic arbor have more spines when compared to cells with unidirectional apical dendritic structure, raising the question of whether synaptic strength differs between these two neuronal subtypes. Therefore, we hypothesized that a reciprocal relationship between synaptic density and strength could be at play between bifurcated and unidirectional CA1 neurons. To test this hypothesis, we next quantified the intensity values of PSD95. Of note, the size of PSDs is strongly associated with synaptic strength, and it has been previously demonstrated that there is a linear, proportional relationship between eGFP-PSD95 brightness (fluorescence intensity) and PSD dimensions.^29,32^ Our results showed a significant increase in fluorescence intensity of PSD95 puncta in both spine and shaft PSDs of unidirectional neurons compared to bifurcated neurons. However, this was not statistically significant when comparing the intensity values in relation to different dendritic compartments, despite the trend being the same, suggesting that this effect was overall cell-type dependent (**Fig. 1E+F**). This indicated that a higher synaptic density correlates with lower fluorescence intensity as observed in bifurcated neurons, and the reverse was found in unidirectional neurons.

### Protein expression profile of shaft and spine PSDs in dissociated hippocampal neurons

Glutamatergic spine synapses have been extensively investigated among different developmental phases, with a thorough characterization of the protein composition in both pre- and postsynaptic compartments.^3,28,33–36^ However, a comprehensive protein expression profile of glutamatergic shaft synapses in mature neurons is thus far missing.

To deepen our understanding, a systematic analysis targeting classical postsynaptic proteins known to be enriched in glutamatergic PSDs was performed in primary rat hippocampal neuron cultures. DIV15 cells were co-transfected with mRuby3 (cell fill) (**Fig. 2A**, left panel) and PSD95.FingR-GFP (postsynaptic excitatory marker) (**Fig. 2A**, middle panel). As the PSD95.FingR-GFP probe enables us to observe endogenous localization of the PSD95 protein, the combination with immunolabeling allowed the colocalization analysis of endogenous PSD95 with the glutamatergic postsynaptic marker of interest in both spines and dendritic shafts (**Fig. 2B**). We then determined the frequency distribution of selected postsynaptic proteins within dendritic spines (**Fig. 2C**) and shafts (**Fig. 2D**) as well as their colocalization with the postsynaptic excitatory marker, PSD95 (**Fig. 2E**). We found that, in basal conditions, scaffold proteins such as Homer, Shank3 and PSD95, as well as SynGAP and cortactin, are found in most spines. Synaptopodin, as a marker of the spine apparatus (SA), was only found to be present in ∼20-30% of the spines, consistent with previous studies.^37,38^ The remaining protein targets Shank1, Shank2, Neuroligin, Ca_V_1.2, AMPAR, NMDAR, and Grip1 were found in 25-50 % of the spines (**Fig. 2C**). All of these proteins were also found enriched in distinct puncta in the dendritic shaft (**Fig. 2B+D**). The analysis revealed that PSD scaffold proteins such as Homer, Shank1, Shank2, Shank3 as well as synaptopodin and Ca_V_1.2 voltage-gated calcium channel colocalize with PSD95 more significantly in spine PSDs than in shaft PSDs (Homer in spine PSDs: ∼81%, in shaft PSDs: ∼32%; Shank1 in spine PSDs: ∼60%, in shaft PSDs: ∼34%; Shank2 in spine PSDs: ∼82%, in shaft PSDs: ∼42%; Shank3 in spine PSDs: ∼83%, in shaft PSDs ∼52%; synaptopodin in spine PSDs: ∼28%, in shaft PSDs: ∼12%, Ca_V_1.2 in spine PSDs: ∼23%, in shaft PSDs: ∼9%) (**Fig. 2E**). In contrast, the other postsynaptic targets, Neuroligin, SynGAP, AMPAR, NMDAR, cortactin and Grip1 demonstrated a similar level of co-occurrence with PSD95 across both spine and shaft PSDs (**Fig. 2E**).

**Figure 2:**
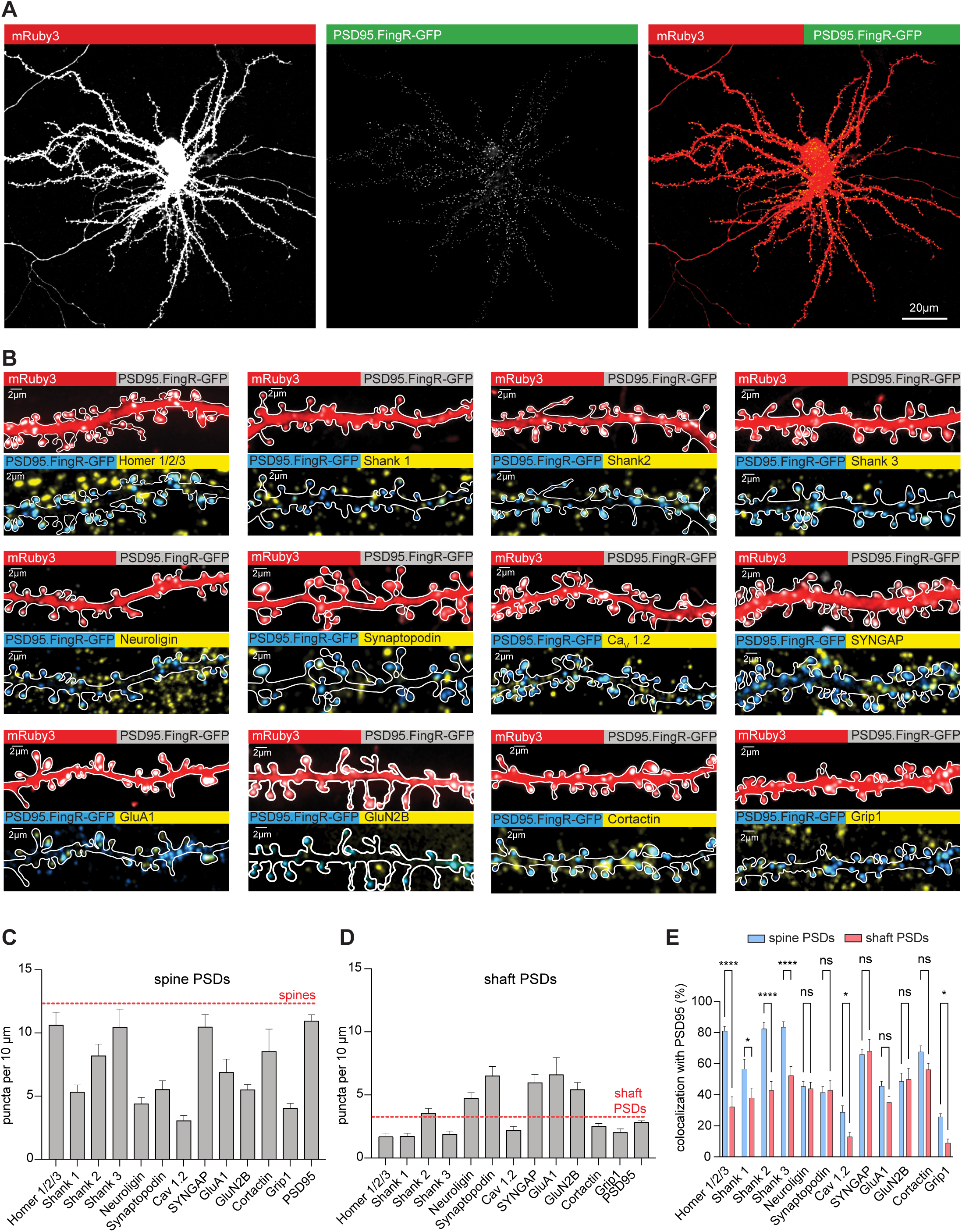
Comparative analysis of spine versus shaft PSD components. **A)** Representative maximum z-projections confocal images of DIV17 primary rat hippocampal neurons expressing mRuby3 (left panel) and PSD95.FingR-GFP (middle panel) and the overlay of both fluorescent signals (right panel). **B)** Representative maximum z-projections confocal images of dendritic segments of DIV17 primary hippocampal neurons showing immunolabeling of common PSD proteins. **C)** Quantification of protein abundance in spines. The number of spines was counted based on the mRuby3 signal and normalized to a dendritic length of 10 µm. The density of spines was, on average, 12.518 per 10 µm (red dashed line). **D)** Quantification of PSD protein abundance within dendritic shafts. The number of shaft PSD were counted based on the mRuby3 signal combined with the PSD95.FingR-GFP signal and normalized to a dendritic length of 10 µm. The density of the overall shaft PSDs is, on average, 3.1 per 10 µm dendrites (red dashed line). **E)** Quantitative colocalization analysis of endogenous PSD95 with PSD proteins in spine PSDs (blue bar graphs) and shaft PSDs (pink bar graphs). Two-way ANOVA with Bonferroni multiple comparisons test; ****p<0.0001 (Homer); **p=0.0198 (SHANK1); ****p<0.0001 (SHANK2); ****p<0.0001 (SHANK3); *p=0.0175 (synaptopodin); *p=0.0495 (GRIP1). Data are presented as mean±SD. n≥15 cells from 3 independent cultures for each protein.

Altogether, these results are consistent with the fact that in mature hippocampal neurons, glutamatergic PSDs are predominantly formed on spines, with far fewer occurring on the dendritic shafts. Overall, the shaft PSDs are similar to spine PSDs regarding their protein composition. The biggest differences were found in the reduced presence of scaffolding proteins in shaft PSDs. Among the tested candidates, no protein could be identified that exclusively targets shaft PSDs.

### Shaft PSDs are coupled to active presynaptic compartments

Next, we asked if shaft PSDs are active synapses. For this, we employed the syt-1 antibody uptake assay in mature rat primary hippocampal neurons (**Fig. 3A**).^8,9^ At DIV17, cells, co-transfected one day earlier with mRuby3 and PSD95.FingR-GFP plasmids were incubated with a syt-1 antibody for 30 minutes, then fixed and imaged. We found that both spine and shaft PSDs colocalize with uptake of syt-1 antibody, as shown by the intensity profiles measured from maximum z-projection images (**Fig. 3B**), suggesting that these are functional synapses. In line with this, approximately 60% of all detected spine PSDs were also positive for syt-1 in spines. Their colocalization significantly decreased to ∼40% when the syt-1 antibody was applied to tetrodotoxin (TTX)-silenced cultures (**Fig. 3C**). About 40% of shaft PSDs were positive for syt-1 under basal conditions. Colocalization significantly decreased to ∼20% after TTX treatment (**Fig. 3C**). These results strongly indicate that both spine and shaft PSDs are opposite of active presynaptic terminals. Furthermore, both spine and shaft synapses show reduced colocalization with internalized syt-1 antibody, highlighting that both spine and shaft PSDs are sites of active synaptic transmission.

**Figure 3:**
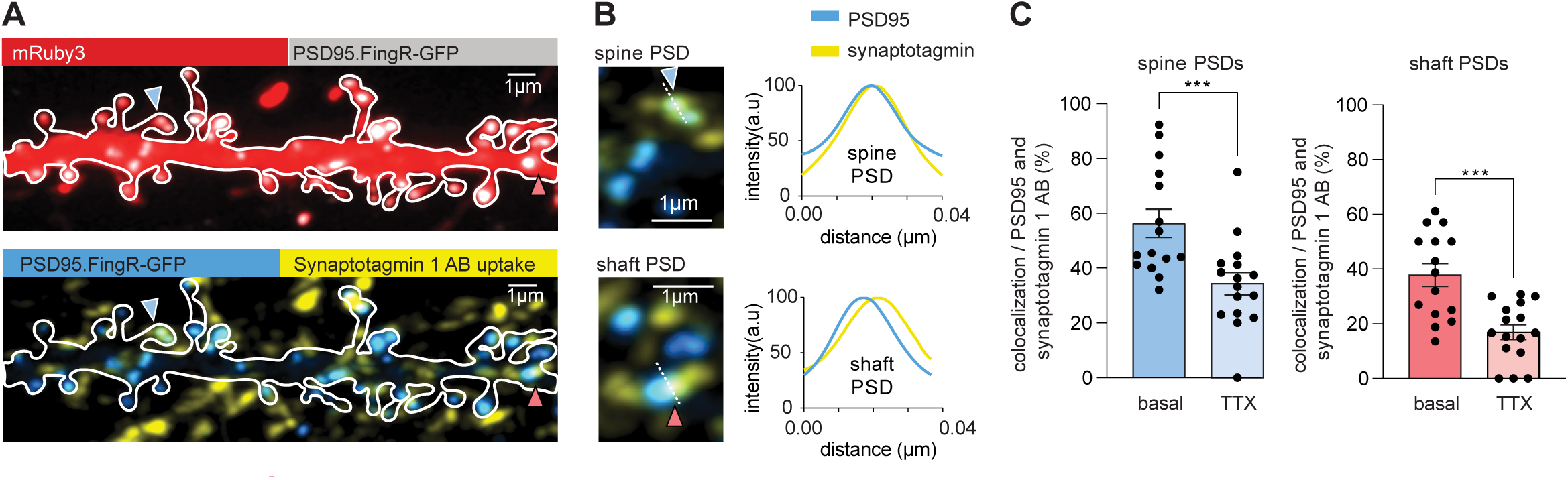
Excitatory shaft synapses are active. **A)** Representative maximum z-projections of confocal images of a dendritic segment of a DIV17 primary hippocampal neuron expressing mRuby3, PSD95.FingR-GFP (top panel) and immunolabeled against syt-1 after live uptake of a syt-1 antibody (AB) by active presynaptic terminals (bottom panel). Example spine PSD marked by the blue arrow and shaft PSD by the pink arrow. **B)** Fluorescence intensity line profile of endogenous PSD95 and syt-1 AB signals at both spine and shaft PSDs. **C)** Silencing of neuronal activity by TTX application reduces presynaptic vesicle release at both spine and shaft PSDs. Mann-Whitney Test. ***p=0.0009 (spine PSDs). ***p=0.0008 (shaft PSDs). Data are presented as mean±SD. All the quantifications have been performed on 15 neurons from 3 independent preparations.

### Shaft synapses undergo structural plasticity

Next, we assessed plasticity-related changes of shaft PSDs in response to stimulations that induce long-term synaptic modifications. It has been shown that there is a dynamic remodeling of postsynaptic organization and composition that usually occurs in response to multiple synaptic inputs and learning.^24,39–45^ Previous studies have shown that in non-stimulated primary hippocampal neurons, shaft PSDs can persist for long periods and rarely transform into spine PSDs.^5^ However, in mouse cortical layer 2/3 pyramidal neurons, repetitive glutamate stimulation alone is sufficient to trigger *de novo* spine growth from dendritic shafts in a very specific manner, a mechanism which is NMDA-receptor dependent.^46^

Our results show that most synaptic proteins expressed in dendritic spines are also located at the shaft PSDs. We hypothesized that possible conversions from shaft to spine PSDs could represent a way for neurons to rapidly respond to long-term plasticity changes, such as in LTP or LTD conditions. To analyze LTP, we performed live-cell imaging using primary rat hippocampal neuron cultures at DIV17-18 before and after glycine-induced cLTP. Cells were co-transfected with PSD95.FingR-GFP and mRuby3. Transfected neurons were imaged for ∼30 min under basal conditions, and for up to 5 h after short-term application of a stimulation solution (**Fig. S2B**).^6,47,48^ The analysis showed that a large fraction of the PSD95 puncta remained stable over time at the synapse locations (indicated by pink arrow in **Fig. 4B**). However, a smaller fraction of puncta was shifting their position within a micrometer range, possibly transported or diffusing along the dendritic shaft, sometimes clustering, and sometimes relocating at the base of the spines. Moreover, we observed a widespread remodeling of dendritic spines upon cLTP induction. We noticed structural membrane changes at shaft PSDs and observed spine-like structures protruding from existing shaft PSDs (indicated by blue arrows in **Fig. 4B**). The quantitative analysis revealed that approximately 6 % of the total detected shaft PSDs transform into spine PSDs with PSD95 relocating to the spine head, whereas approximately 20 % of the shaft PSDs shift position along the dendritic segment. The largest proportion, approximately 70 % of shaft PSDs, showed high stability and did not exhibit any changes in terms of dendritic location (**Fig. 4C**).

**Figure 4:**
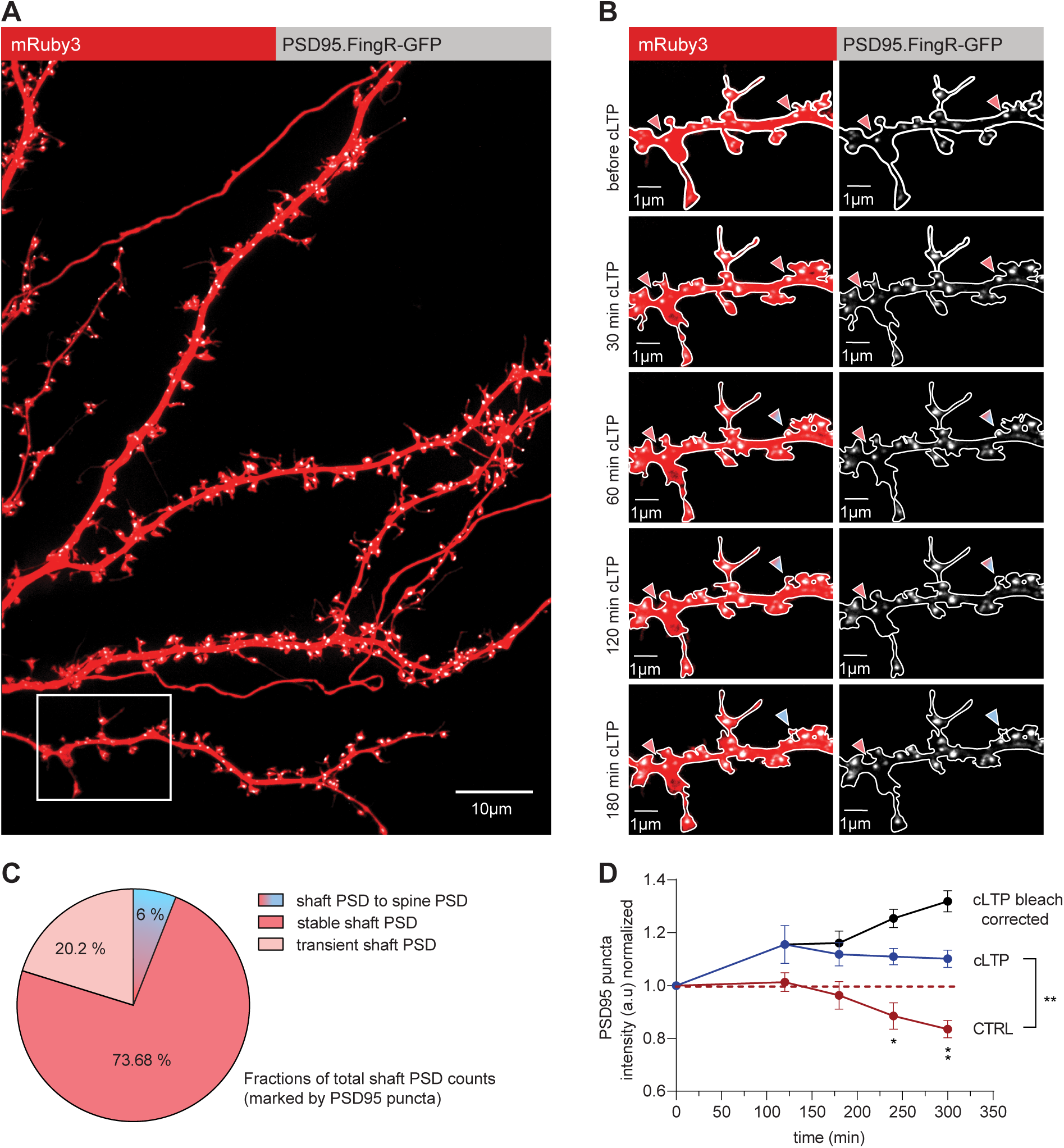
Most shaft PSDs are stable over extended time periods, with some transient shaft PSDs and a subset of shaft PSDs turning into spine PSDs upon cLTP. **A)** Representative maximum z-projections of confocal images of a DIV17 primary rat hippocampal dissociated neuron expressing mRuby3 and PSD95.FingR-GFP. **B)** Dendritic segment of the transfected neuron imaged before and after cLTP induction, showing stable shaft PSDs indicated with pink arrows and shaft PSDs transforming into spine PSDs indicated with pink to blue transition arrows. **C)** Quantification of the proportion of shaft PSDs turning into spine PSDs, stable shaft PSDs, and transient shaft PSDs. **D)** Normalized intensity of endogenous PSD95 signal after cLTP induction. PSD95 signal increases for about 5 h (blue curve) upon 5 min application of the cLTP induction solution. On the contrary, in basal condition (CTRL) neurons show a decrease in PSD95 intensity over a 5 h timeframe (red curve), suggesting photobleaching. The amount of photobleaching detected without stimulation was used to correct the PSD95 signal after cLTP induction (black curve). Two-way ANOVA with Šídák’s multiple comparisons test; **p=0.0031. Quantification of cLTP was performed in 10 neurons from 4 independent preparations. Quantification of basal condition (CTRL) was performed in 6 neurons from 1 preparation. Data are presented as mean±SEM.

An increase in the PSD size is fundamental for LTP maintenance.^39,49–51^ Therefore, we estimated the level of PSD95 expression in shaft and spine PSDs before and after cLTP. As expected, the quantification showed a significant increase in expression over time in response to cLTP (**Fig. 4D**, blue curve), but not under basal conditions (**Fig. 4D**, dark red curve).

In summary, these results provide new key insights into the plasticity-related changes of excitatory shaft synapse structures and highlight the potential for pre-existing shaft synapses to transform into dendritic spine synapses under certain activity conditions. Importantly, most of the shaft PSDs are remarkably stable over time, indicating that they represent an independent synapse type that participates in neurotransmission.

### Long-term depression of excitatory synapses

Next, we aimed to investigate the impact of LTD on the number of shaft and spine PSDs.^37^ We employed a protocol for NMDA receptor-dependent LTD (cLTD) in mature primary rat hippocampal neurons transfected with mRuby3 (cell fill) and PSD95.FingRs-GFP and immunolabeled for Bassoon (**Fig. 5A**). cLTD was induced by a brief (3 min) application of NMDA (30 µM)^23^, and we assessed the synaptic density of both spine and shaft PSDs at two time points: under basal conditions and 3 h after induction of LTD.

**Figure 5:**
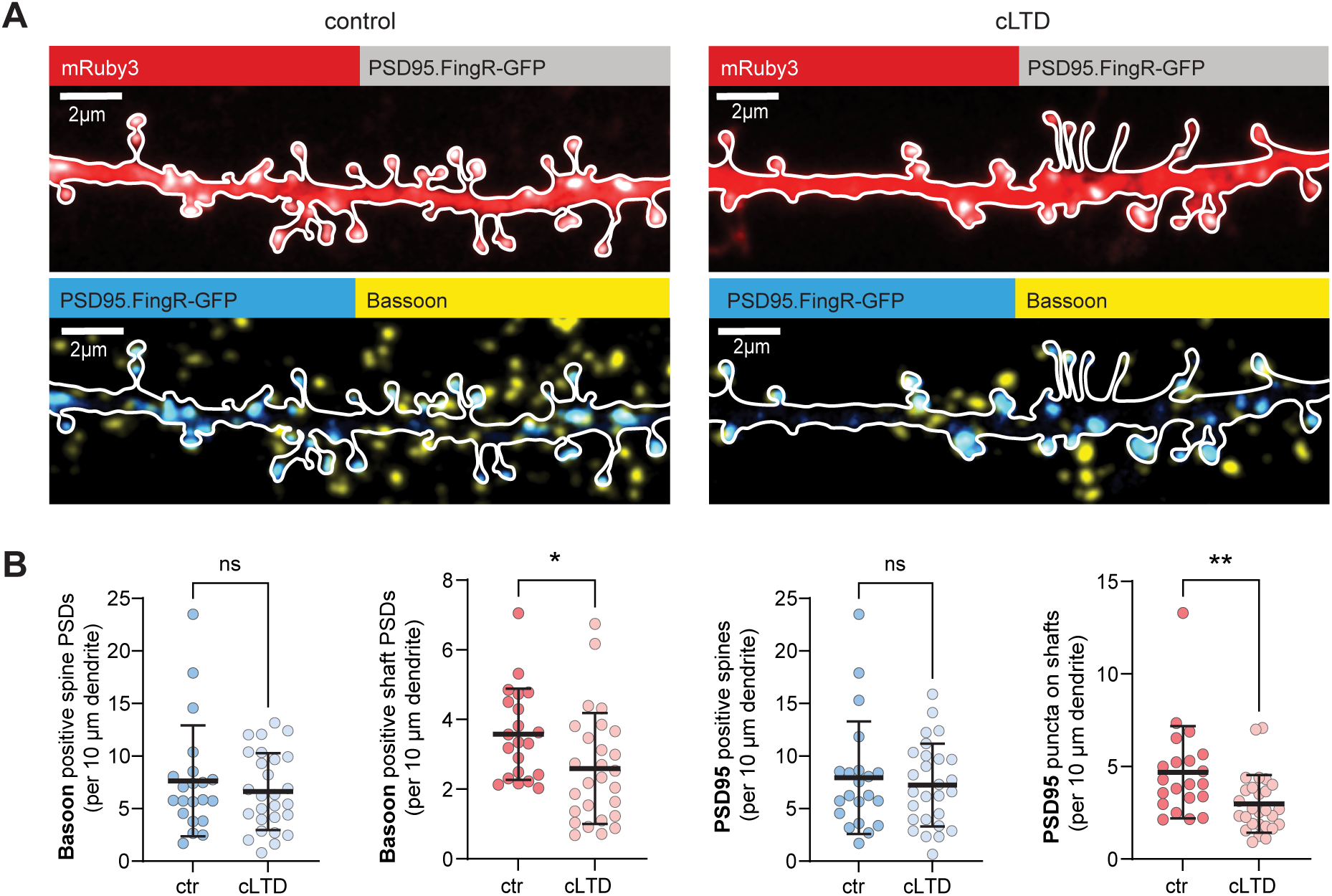
Shaft synapses are pruned upon cLTD. **A)** Representative maximum z-projections of confocal images of dendritic segments of DIV17 primary rat hippocampal neurons expressing mRuby3, PSD95.FingR-GFP and immunolabeled for Bassoon before and after NMDA-mediated cLTD. **B)** Quantification of synapse densities calculated by counting the number of spine and shaft PSDs positive for Bassoon and PSD95. Unpaired non-parametric two-tailed t-test; **p=0.0057 PSD95 puncta on shafts; *p=0.0268 Bassoon positive shaft PSDs. Data are represented as mean±SD.

Our findings indicate that while dendritic spines maintained a consistent density of synapses positive for the presynaptic marker Bassoon, there was a significant decrease in the number of synaptic contacts on shaft PSDs post-LTD (**Fig. 5B**), suggesting that LTD preferentially affects synaptic contacts on shaft PSDs compared to spine PSDs. This result was further corroborated by similar observations using the postsynaptic marker PSD95, where we found that while the density of spine PSDs remained unchanged, there was a significant reduction in shaft PSDs following LTD (**Fig. 5B**). These results suggest that synapses located on dendritic shafts may be more vulnerable to modifications induced by NMDA-dependent LTD than those on dendritic spines.

## Discussion

In this study, we performed quantitative analysis of synaptic distribution in mature CA1 glutamatergic neurons and could show that, in addition to the spinous localization, around 25 % of excitatory postsynaptic sites are located on the dendritic shafts. This distribution is independent of the overall structure of the dendritic arbor in two classes of neurons: unidirectional and bifurcated. The shaft PSDs are highly similar to spine PSDs regarding general protein content and architecture. However, there are subtle differences between spine and shaft PSDs in the relative abundances of scaffold proteins. Like spines, shaft PSDs have adjacent presynaptic sites and participate in ongoing synaptic activity. Upon induction of cLTP, the size of shaft PSDs increases, but most shaft PSDs remain on the shaft with only a small fraction turning into spines. We found that during cLTD, synaptic loss occurs mostly at the expense of shaft synapses.

Using high-resolution confocal microscopy, we were able to quantify the abundance and location of PSD95 puncta throughout whole CA1 neurons. Organotypic hippocampal slice culture preserves the native circuitry and dendritic architecture.^52–55^ This makes them particularly well-suited to dissect the shaft and spine localization of excitatory synapses in a near-physiological context. Overall, we found that bifurcated neurons have a higher density of PSDs, but their intensities are weaker. The abundance of PSD scaffolds may be a limiting factor, thereby the number and size of PSDs are counterbalancing each other. On a larger scale, this could be a mechanism to regulate excitability by the expression levels of key PSD proteins.

The characterization of the protein composition of shaft PSDs in dissociated primary rat hippocampal neuron culture showed that they share the core architecture of excitatory postsynapses with their spine PSD counterparts. This includes glutamate receptors, adhesion molecules, Ca^2+^ channels, and scaffold proteins like PSD95, Shanks, and Homer. The similar composition, albeit with reduced scaffold protein expression in shaft PSDs, is in line with the literature on shaft PSDs, although no characterization of the shaft PSDs in glutamatergic neurons has been performed to date.^56,57^ In the future, it remains to be tested whether any proteins are exclusively present in the shaft or spine synapses. The demonstration that shaft synapses are sites of active synaptic transmission, together with their expression profile containing AMPARs and NMDARs, supports their classification as *bona fide* synapses rather than immature or silent synapses. Super-resolution microscopy, fluorescence recovery after photobleaching (FRAP) experiments, and single-molecule tracking of shaft PSD components could reveal the nanostructure of these synapse types and the dynamics and exchange of their molecular composition. Due to their localization on the dendritic shafts, they can circumvent the electrical and biochemical compartmentalization of spines, thereby influencing the excitability of dendrites and the spreading of dendritic Ca^2+^ signals.^6,58,59^ Future research could address the contribution of shaft synapses to the circuits as well as their impact on dendritic compartmentalization.

By applying established cLTP and cLTD protocols, we were able to show that shaft synapses are plastic. Using long-term live imaging, we found that cLTP causes an increase in PSD95 puncta intensity, which indicates PSD enlargement. Additionally, shaft synapses were remarkably stable over extended timespans. A small proportion of shaft PSDs could transform into spine PSDs. This process is likely F-actin dependent, as it has been shown that shaft synapses are surrounded by a dense F-actin mesh and remodeling of spines is F-actin dependent.^8,18^ Future modeling studies could estimate the contribution of these newly added spines to neuronal excitability and function. In contrast to cLTP, the cLTD paradigm induced synaptic weakening within minutes, but the structural synapse loss typically emerges more slowly, often requiring several hours to manifest.^60,62^ Our 3 h cLTD protocol was sufficient to reduce the number of shaft synapses, indicating that these contacts are particularly vulnerable to LTD-associated destabilization. In contrast, dendritic spines remained structurally intact at this time point, suggesting that spine elimination may follow a slower or mechanistically distinct process compared with shaft synapse loss. The relative resistance of spine synapses to LTD in our study is in line with previous work, indicating that spines may possess distinct molecular machinery or signaling pathways that confer greater resilience to LTD-induced alterations.^61,65^ It is possible that the lower abundance of the scaffold proteins Shanks and Homer leads to reduced stability of shaft PSDs.

The specific location of postsynaptic sites directly on the dendritic shaft allows for different interactions with other cellular organelles. Especially, vesicular organelles or mRNA granules can be positioned in proximity to the shaft PSDs. Earlier work shows that F-actin patches in dendrites and enrichment of F-actin at presynaptic boutons enable vesicle stalling by myosin motor proteins.^8,63^ Thereby, shaft PSDs could serve as hubs for secretory trafficking organelles to interact and mediate their function. The activity of processive myosins, which are involved in organelle stalling, is regulated by Ca^2+^ and calmodulin.^37,63^ Synaptic activity could therefore locally cause organelle stalling and enable interaction of different organelles to allow for compartmentalized protein and membrane turnover. For instance, mTORC1, a central regulator of cellular metabolism and protein synthesis, is activated at the lysosomal surface.^68,70^ Thus, stalling of lysosomes at shaft synapses could provide spatial control for lysosome-dependent Ca²⁺ signaling and enable localized mTORC1 activation. Such positioning would create dendritic compartments where lysosomes, translation machinery, and signaling complexes intersect, thereby supporting highly localized regulation of protein synthesis and synaptic remodeling.

As dendritic spine formation and removal have been shown to be highly relevant for learning and memory formation^24^, future work on excitatory shaft synapses could focus on their role in these processes, as they have been shown to be highly plastic synapses with great potential to be potentiated and easy to be pruned.

## Materials and Methods

### Animals

Wistar Crl:WI (Han; Charles River), Wistar Unilever HsdCpb:WU (Envigo), and Wistar RjHan:WI (Janvier Labs) rats were used in this study. Pregnant rats (E18) for primary hippocampal cultures and male rat pups aged between P4-P7 for organotypic hippocampal slice cultures were sacrificed in accordance with the Animal Welfare Law of the Federal Republic of Germany (Tierschutzgesetz der Bundesrepublik Deutschland, TierSchG) and with the approval of local authorities of the city-state Hamburg (Behörde für Gesundheit und Verbraucherschutz, Fachbereich Veterinärwesen, from 21.04.2015), the animal care committee of the University Medical Center Hamburg-Eppendorf (permit numbers ORG781 and 1035), and the animal welfare committee of the Charité (permit number T-CH 0002/21).

### Organotypic slice cultures and single-cell electroporation

Organotypic hippocampal slices were prepared from male Wistar rats at post-natal day 4-7 as described previously.^55^ Briefly, dissected hippocampi were cut into 400 μm slices with a tissue chopper (McIlwain Tissue Chopper; Model TC752) and cultured on a hydrophilic polytetrafluoroethylene (PTFE) membrane (Millicell CM, Millipore). Cultures were maintained at 37 °C, 5 % CO_2_ in a medium containing 78.8 % MEM, 20 % heat-inactivated horse serum supplemented with 1 mM L-glutamine, 0.00125 % ascorbic acid, 0.01 mg/ml insulin, 1 mM CaCl_2_, 2 mM MgSO_4_, 13 mM D-glucose, and 20 mM HEPES. No antibiotics were added to the culture medium. Individual CA1 pyramidal cells were transfected by single-cell electroporation^54,64^ using pAAV-synapsin-mRuby3 and PSD95.FingR_eGFP_CCR5TC plasmids at a final concentration of 1 ng/µL. During electroporation, slices were perfused with a solution containing: 145 mM NaCl, 10 mM HEPES, 12.5 mM D-glucose, 1.2 mM NaH_2_PO_4_, 2.5 mM KCl, 1 mM MgCl_2_, and 2 mM CaCl_2_. The pH was adjusted to 7.4. 2-4 days after transfection, electroporated slices were fixed with 4 % Roti-Histofix/4 % sucrose for 15 min at room temperature (RT), washed thrice for 10 min each with phosphate-buffered saline (PBS), and mounted on microscope slides with Mowiol (Carl Roth) mounting media. Mowiol was prepared according to the protocol provided by the manufacturer.

### Primary hippocampal neuron preparation and transfections

Primary rat hippocampal neuron cultures were prepared as described previously with minor adjustments.^8^ In brief, hippocampi were isolated from E18 embryos in ice-cold HBSS and mechanically dissociated after 10 min of 0.25 % trypsin treatment at 37 °C, then plated on poly-L-lysine-coated glass coverslips (18mm) at a density of 40.000-60.000 cells/mL suspended in DMEM supplemented with 10 % fetal bovine serum and 1x penicillin/streptomycin antibiotic. The plating medium was replaced after 1 h by BrainPhys neuronal medium supplemented with SM1 and 0.5 mM glutamine. Primary neurons were grown in an incubator at 37 °C, 5 % CO_2_, and 95 % humidity. Cells were transfected at DIV15 by using Lipofectamine 2000 and a total DNA/lipofectamine ratio of 1:2. Before transfection, the original neuronal medium was removed and kept at 37 °C. BrainPhys medium without additional supplements was added to the cells for transfection. To this medium, the pre-mixed and pre-incubated transfection mix consisting of plasmid DNA, lipofectamine 2000, and BrainPhys, was added. After 60-75 min, the transfection medium was exchanged back to the original BrainPhys containing supplements. The pAAV-synapsin-mRuby3 and pCAG_PSD95.FingR-eGFP-CCR5TC plasmids were transfected at a concentration of 1 µg/well. 2 days after the transfection, the experiments were conducted.

### Chemical LTP induction

For chemical induction of LTP, DIV17-21 primary rat hippocampal neurons grown on coverslips were stimulated with a glycine-based solution (**Fig. S2B**). Briefly, coverslips containing neurons were moved to a microscopy chamber and perfused with aCSF (Mg^2+^-free) containing: 145 mM NaCl, 10 mM HEPES, 12.5 mM D-glucose, 1.2 mM NaH_2_PO_4_, 2.5 mM KCl, and 2 mM CaCl_2_, pH adjusted to 7.4. Then, cells were stimulated by perfusing aCSF (Mg^2+^-free) solution containing 200 µM glycine, 1 µM TTX, and 50 µM 1(S),9(R)-(−)-Bicucullin-Methiodide for 5 min.

### Chemical LTD induction

For chemical induction of LTD, DIV17-21 primary hippocampal neurons were stimulated with NMDA. Briefly, 30 µM NMDA was prepared in BrainPhys medium and pre-warmed to 37 °C. Conditioned medium from the cells was removed and collected separately at 37 °C, while neurons were incubated with NMDA-containing medium for 3 min. Subsequently, the NMDA medium was replaced with the previously collected conditioned medium, and neurons were moved back to the incubator for 3 h. Finally, cells were fixed with 4 % Roti-Histofix / 4 % sucrose and then used for immunocytochemistry.

### Immunocytochemistry (ICC)

Dissociated primary neurons were fixed with 4 % Roti-Histofix / 4 % sucrose for 20 min at room temperature (RT), washed thrice for 10 min each with PBS, and permeabilized with 0.2 % Triton X-100 in PBS for 15 min at RT. The neurons were then washed thrice in PBS and blocked for 60 minutes at RT with blocking buffer (10 % horse serum, 0.1 % Triton X-100 in PBS). Primary antibodies (**Table 3**) were diluted in blocking buffer and added to the cells at 4°C overnight. Subsequently, the neurons were washed thrice for 10 min each, and secondary antibodies (**Table 4**) were diluted in blocking buffer, added to the cells, and incubated for 1 h at RT. Finally, cells were washed thrice for 10 min each in PBS and mounted on microscopy slides with Mowiol. Mowiol (Carl Roth) was prepared according to the protocol provided by the manufacturer.

**Table 1:**
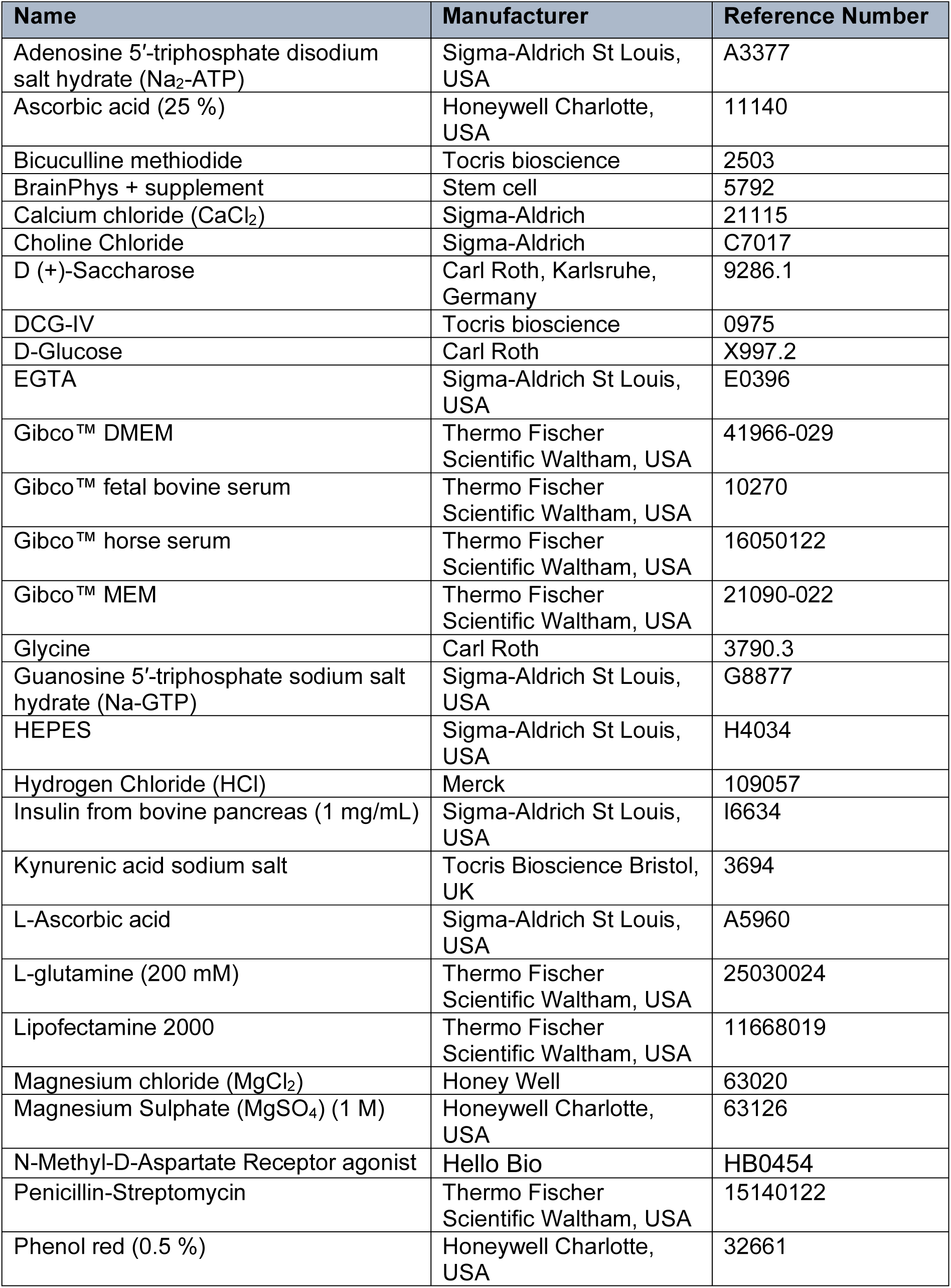
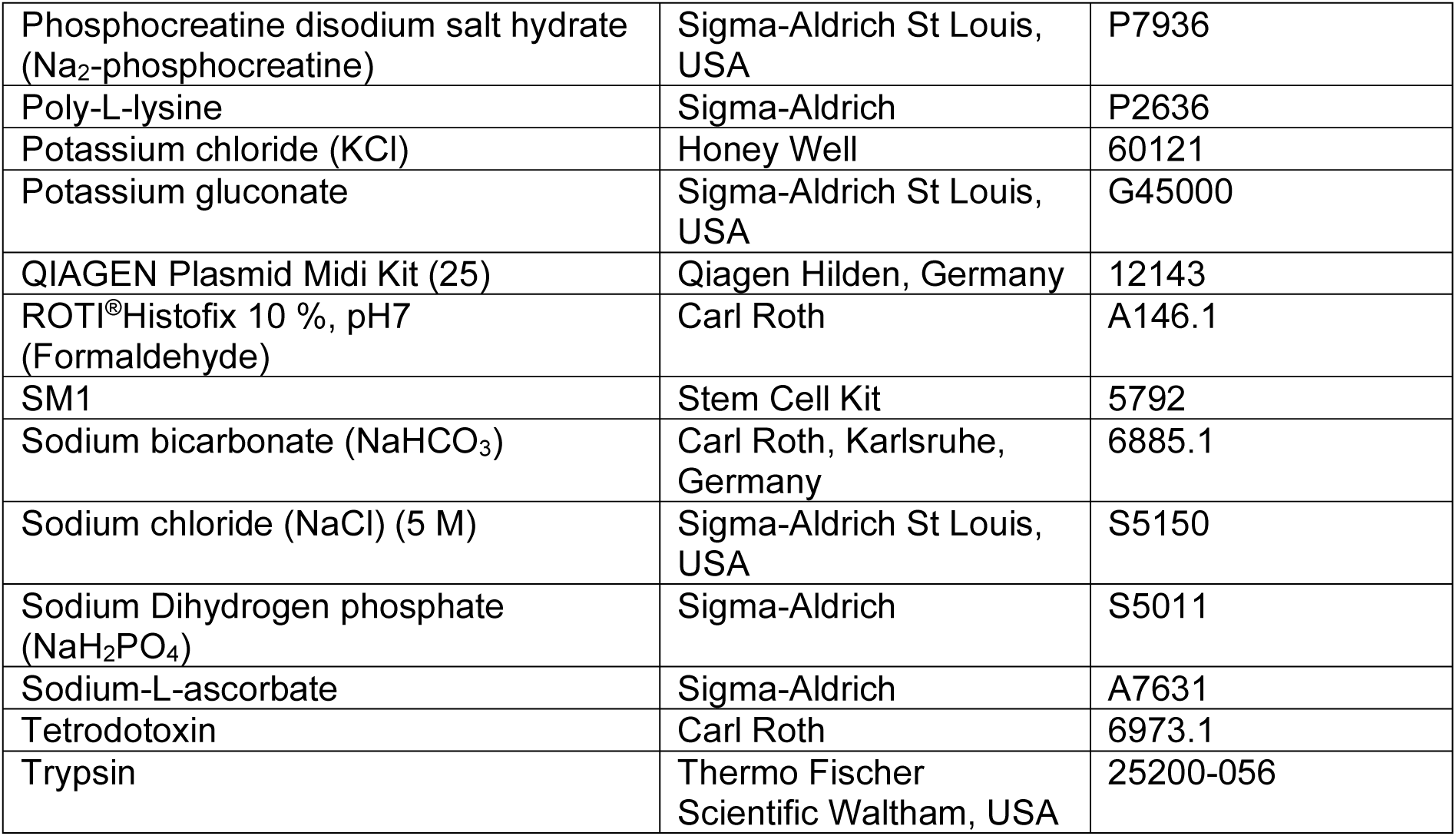
Chemicals/Reagents.

**Table 2:** Plasmids.

| Name | Source | Reference Number |
| --- | --- | --- |
| pAAV-synapsin-mRuby3 | Gift by Bas van Bommel | N/A |
| pCAG_PSD95.FingR-eGFP CCR5TC | Addgene | 46295 |

**Table 3:**
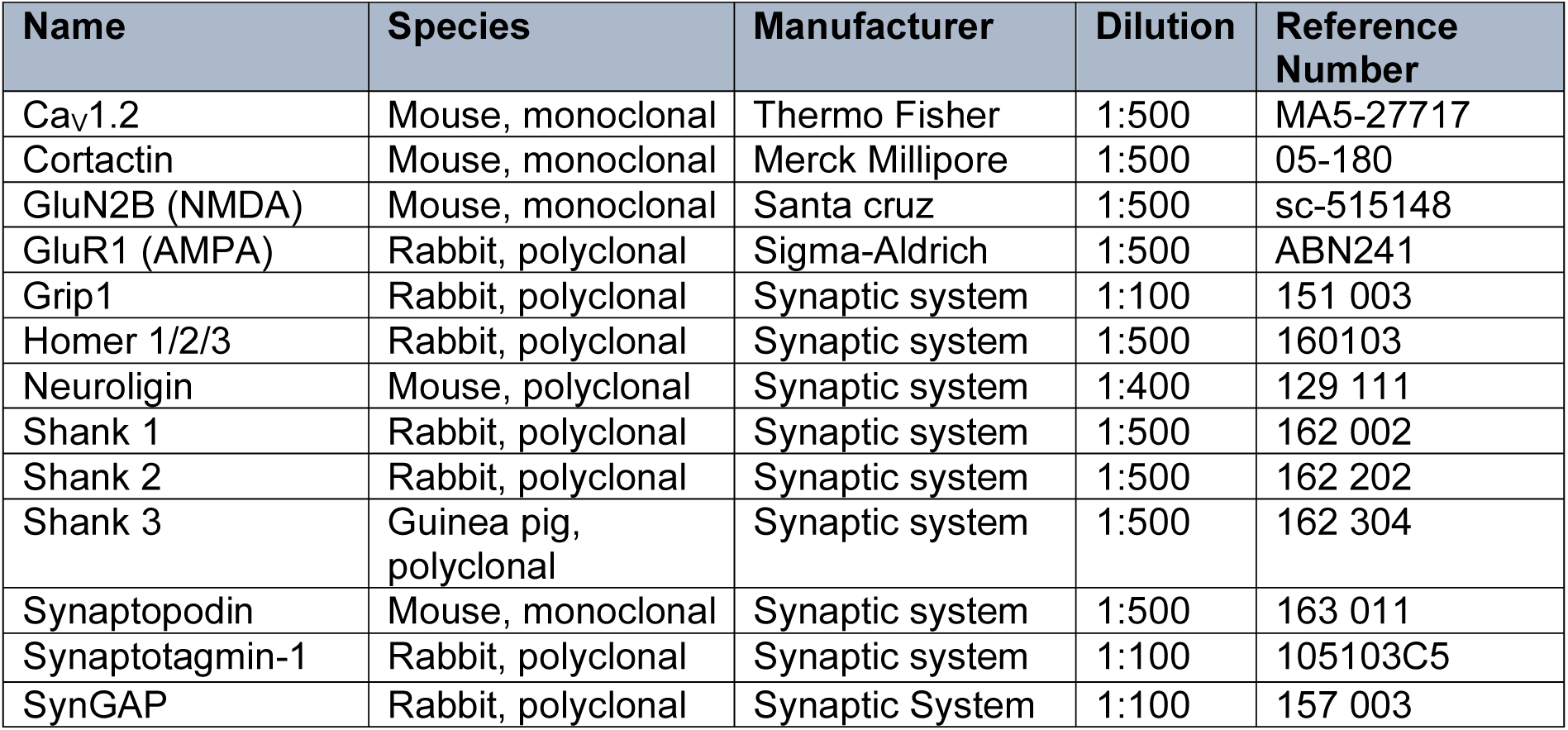
Primary antibodies.

**Table 4:** Secondary antibodies.

| Name | Species | Manufacturer | Dilution | Reference Number |
| --- | --- | --- | --- | --- |
| Alexa 647 | Rabbit, polyclonal | Life Technologies | 1:500 | A21245 |
| Alexa 647 | Mouse, polyclonal | Life Technologies | 1:500 | A21235 |
| Alexa 635 | Guinea pig, polyclonal | Abberior | 1:500 | st635p |

**Table 5:** Software.

| Name | Developer | Application |
| --- | --- | --- |
| Adobe Illustrator | Adobe Inc. San Jose, California | Figures |
| GraphPad Prism | GraphPad, Boston | Graphs, Statistics |
| Image J | National Institute of Health Bethesda, USA | Imaging Analysis |
| Imaris microscopy image analysis software | Oxford Instruments Abingdon, UK | 3D reconstruction |
| Microsoft Office | Microsoft Corporation Washington, US | Word, Excel |
| VisiView | Visitron Systems Puchheim, Germany | Microscopy |

### Fixed cell imaging: spinning disk confocal microscopy

Imaging of fixed samples was performed by using two different confocal microscopy systems. First system: Nikon Eclipse Ti-E inverted microscope controlled by VisiView software (Visitron System). The microscope was equipped with 488, 561, and 647 nm excitation lasers, coupled to a CSU-X1 spinning disc unit (Yokogawa) via a single-mode fiber. Emission light was collected on an Orca flash 4.0LT CMOS camera (Hamamatsu) through a quad-band filter (Chroma, ZET 488/561/647 nm). Z-plane images with a 0.2 µm step size of fixed hippocampal slices and primary neurons were taken with a 100× (Nikon, ApoTIRF 100×/1.49) or a 60× (Nikon, Plan Apo λ 60×/1.40) objective. The second system was provided by the microscopy facility at the German Electron Synchrotron (DESY) center of research. Fixed cell imaging of primary rat hippocampal cultures was performed with a Nikon Eclipse Ti2 inverted microscope controlled by the NIS-Elements software. The microscope was equipped with 488, 561, and 647 nm excitation lasers, coupled to a CSU-W1 spinning disc unit (Yokogawa) via a single-mode fiber. Z-plane images with a 0.2 µm step size of fixed primary neurons were acquired with a 100x (Nikon, CFI Plan Apochromat Lambda 100x/1.45) objective.

### Live-cell Imaging: spinning disk confocal microscopy

Live-cell imaging was performed with the same spinning disk microscopy systems described in the previous paragraph. Live imaging of primary neurons was performed for up to 6 h by using a Ludin chamber (Life Imaging Services) (**Fig. S2B**) and µ-Dish 35 mm, or high Glass Bottom (Ibidi). The focus was maintained by Nikon’s built-in perfect focus system. At both systems, correct temperature (37 °C), CO_2_ (5 %), and humidity (90 %) were maintained with a stage top incubator (Okolab). Pharmacological treatments were performed during live imaging by adding and washing out the drugs by gently pipetting in and out of the recording chamber. A set of z-stack images was acquired for 30 min (1 z-stack every 10 min) before cLTP induction. After stimulation, the neurons were placed back in the original Mg^2+^-free aCSF, and live imaging was performed up to 5 h (1 z-stack every 30 min).

### Distinction of dendritic compartment and main apical branching

Maximum projections and z-planes, utilized for the analysis, were processed and evaluated using ImageJ software.^66^ The analysis aimed to quantify the total number of PSD95-positive spines and shaft PSDs within different dendritic compartments of two morphologically distinct CA1 hippocampal neuronal types as classified by Benavides-Piccione and colleagues.^30^ In this study, the authors did not provide a precise definition of the main apical branching; therefore, an initial classification was devised. CA1 cells were classified as bifurcated if a branching point was observed in the main apical shaft, within approximately 200 µm away from the soma, producing two continuing shafts of approximately equal diameter post bifurcation. CA1 cells in which no branching point was observed, or where a branching point produced two morphologically different shafts, were classified as unidirectional. Synaptic density was quantified for unidirectional neurons (no bifurcation of main apical shaft) compared to bifurcated neurons (singular bifurcation of main apical shaft). The distinguished dendritic compartments were the following: main apical shaft, basal dendrites, tuft dendrites, and oblique dendrites. Only primary and secondary branches were considered for analysis.

### Identification of spine and shaft PSDs

Spines were identified as mRuby3-positive protrusions extending from the dendritic body. Spines of different morphologies (filopodia, mushroom, stubby, etc.) were included in the analysis. PSD95 was used as a postsynaptic marker and was detected with PSD95.FingR-GFP, a probe based on an intrabody against PSD95 fused to a GFP.^26^ PSD95.FingR-GFP reports the localization and amount of the endogenous target protein. Spines were defined as “protrusions” or “PSD95-positive” depending on the presence of PSD95. Synapses appearing above or below the focal planes of the dendritic shaft were identified as protrusions growing in the z dimension and therefore not considered as shaft synapses. To confirm this spine identification method, 3D reconstructions of representative dendritic segments were constructed using the filament tracer tool in Imaris microscopy image analysis software (**Fig. S2A**). This method allowed for the differentiation of protrusions and PSD95-positive shaft PSDs. Shaft PSDs were identified as puncta of PSD95 signal within the dendritic shaft. Spine and shaft PSDs were counted manually using the ImageJ cell counter tool and then normalized to a dendritic length of 10 µm. In every neuron, three separate dendritic segments (15 - 60 µm in length) were analyzed for each type of dendrite.

### Three-dimensional reconstruction of dendritic segments

Representative three-dimensional (3D) reconstructions of five dendritic segments were created using the filament tracer tool of Imaris microscopy image analysis software (**Fig. S2A**). z-planes were converted from tif to IMS (Imaris file format) and imported into the software. mRuby3 and PSD95 were selected as two different channels. Scrolling through the individual planes of the mRuby3-positive z-planes, certain circular regions were visible before or after the illumination of the dendritic shaft. As mRuby3 was employed as a morphological marker, we hypothesized that these circular regions were protruding spines perpendicular to the dendritic shaft, illuminated independently of the cell body due to protrusion into a different z-plane. mRuby3 signal was reconstructed in three dimensions with the filament tracer tool using automatic detection and adjusted manually with reference to the Z-plane. PSD95 puncta were detected with the spot detection tool set to identify puncta with a diameter between 0.15 µm to 0.8 µm. The rotation of the 3D reconstructions allowed for the determination of the localization of the PSD95 puncta within the dendritic shaft or on a protrusion. The reconstructions served as a comparative method to determine whether indistinguishable protrusions and PSD95-positive shaft PSDs were correctly identified from z-planes. This method serves as independent validation for the 2D analysis done with ImageJ and additionally contributes to reducing the number of wrongly detected or assigned PSD95-signals. The voxel size was x: 0.065, y: 0.065, and z: 0.2 for all reconstructions.

### Analysis of signal intensity of PSD95.FingR-GFP in hippocampal slices

The intensity of PSD95.FingR-eGFP signal, detecting endogenous PSD95 in spine and shaft PSDs, was determined for five previously analyzed unidirectional and bifurcated neurons. The transcriptional control system of the GFP-fused recombinant probe is matched to the expression of the target protein, therefore, the intensity of PSD95.FingR-GFP fluorescence indicates the amount of PSD95 in an individual PSD ^67^. Three dendritic segments (15-60 µm in length) per dendritic segment type of each neuron were analyzed. Maximum z-plane projections of the 488 nm channel were used as source images for the analysis in ImageJ. Elliptical and polygon selection tools were employed to trace the perimeter of the PSD95 puncta. The integrated density value was determined, and the background intensity was subtracted (available method already used previously).^69^

### Determination of PSD95.FingR-GFP intensity signals in hippocampal primary cultures

The intensity of PSD95.FingR-GFP signal, detecting endogenous PSD95 in the spine and shaft, was determined for 12 primary neurons in total. Dendritic sections of imaged neurons were analyzed using the ComDet plugin of ImageJ, and four independent preparations were quantified. The analysis was performed using maximum intensity z-projections of the 488 nm channel, and integrated density values were determined after background intensity subtraction.

## Acknowledgements

We would like to thank Bas van Bommel for cloning the pAAV-synapsin-mRuby3 construct and for technical assistance with single-cell electroporation. Also, we would like to thank Julia Bär, Maria Andres-Alonso, and Lisa Mallis for the preparation of primary hippocampal neuron cultures. Furthermore, we would like to thank Kay Grünewald and Roland Thünauer for access to the DESY microscopy facility.

## Author contributions

Tomas Fanutza: Conceptualization, Validation, Formal analysis, Investigation, Writing – Original Draft, Visualization, Supervision; Yannes Popp: Validation, Formal analysis, Investigation, Writing – Original Draft, Visualization; Arie Maeve Brueckner: Validation, Formal analysis, Investigation, Writing – Original Draft, Visualization; Nathalie Hertrich: Investigation; Judith von Sivers: Formal analysis; Matthew Larkum: Resources, Funding acquisition; Sarah Shoichet: Resources, Supervision; Marina Mikhaylova: Resources, Supervision, Project administration, Funding acquisition.

## Abbreviations

AB: antibody
aCSF: artificial cerebro-spinal fluid
AMPA: 2-amino-3-(3-hydroxy-5-methyl-isoxazol-4-yl) propanoic acid
AMPAR: 2-amino-3-(3-hydroxy-5-methyl-isoxazol-4-yl) propanoic acid receptor
cLTD: chemical long-term depression
cLTP: chemical long-term potentiation
CTRL: control
DESY: german electron synchrotron
DIV: days in vitro
F-actin: filamentous actin
Fig: Figure
FRAP: fluorescence recovery after photobleaching
LTD: long-term depression
LTP: long-term potentiation
NMDA: N-methyl-D-aspartate
NMDAR: N-methyl-D-aspartate receptor
PBS: phosphate-buffered saline
PSD: postsynaptic density
PTFE: polytetrafluoroethylene
RT: room temperature
SA: spine apparatus
SD: standard deviation
SEM: standard error of the mean
STED: stimulated emission depletion
STORM: stochastic optical reconstruction microscopy
Syt-1: synaptotagmin 1
TTX: tetrodotoxin

---

**Figure S1:**
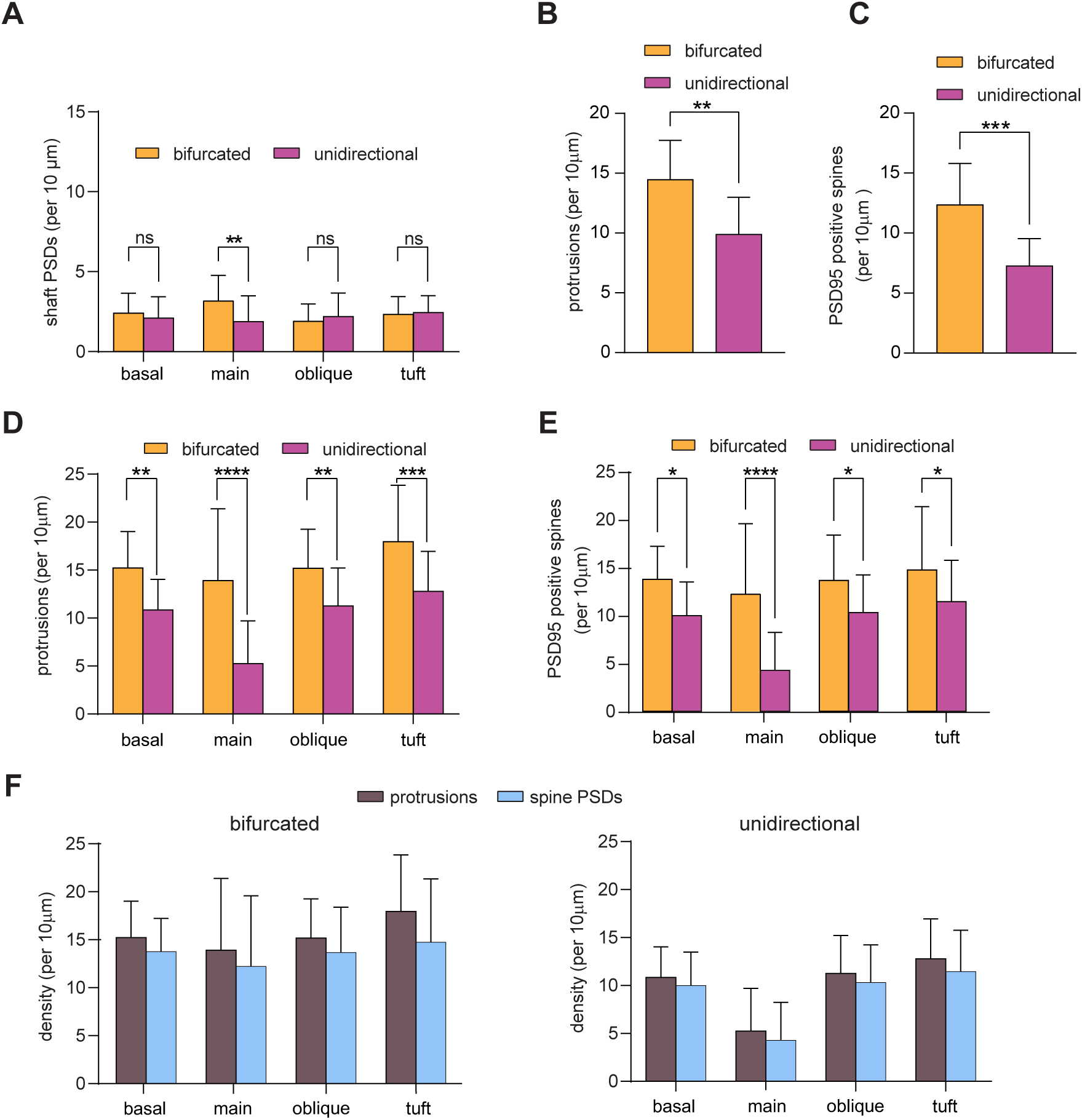
Glutamatergic postsynapses are mainly located on spines and much less within the shaft of mature CA1 pyramidal cell dendrites. **A)** Bar graphs showing the quantification of shaft PSDs in relation to dendritic compartments of bifurcated and unidirectional CA1 neurons. Two-way ANOVA with Šídák’s multiple comparisons test. **p=0.0026 (main dendrite). **B)** Quantification of protrusion density of bifurcated and unidirectional neurons. Mann–Whitney Test. **p=0.0091. **C)** Quantification of overall spines containing PSD95 of bifurcated and unidirectional neurons. Mann–Whitney Test. ***p=0.0008. **D)** Quantification of protrusion density in dendritic compartments of bifurcated and unidirectional neurons. Two-way ANOVA with Šídák’s multiple comparisons test. **p=0.0024 (basal dendrites); ****p<0.0001 (main shaft); **p=0.0080 (oblique dendrites); ***p=0.0002 (tuft dendrites). **E)** Quantification of PSD95-positive spines in dendritic compartments of bifurcated and unidirectional neurons. Two-way ANOVA with Šídák’s multiple comparisons test. *p=0.0139 (basal dendrites); ****p<0.0001 (main dendrite); *p=0.0363 (oblique dendrites); *p=0.0442 (tuft dendrites). **F)** Quantification of the density of protrusions and spine PSDs across dendritic compartments of bifurcated neurons and unidirectional neurons. All data is collected from n=24 dendritic segments from 8 bifurcated neurons. n=33 dendritic segments from 11 unidirectional neurons. Data are presented as mean±SD.

**Figure S2:**
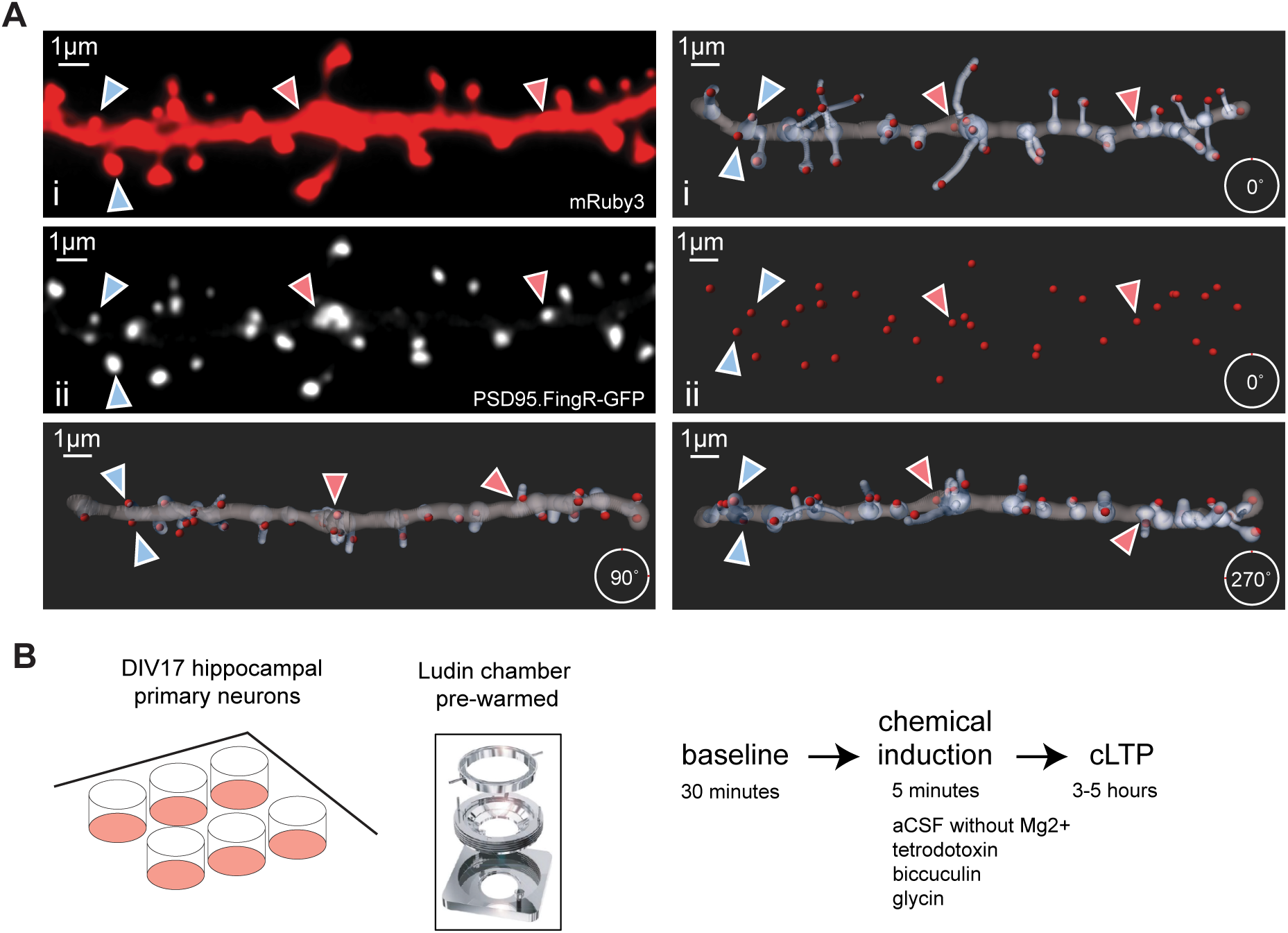
Spine and shaft PSD identification validated by using 3D reconstruction and schematic of cLTP protocol. **A)**Representative maximum z-projection of confocal images of a dendritic segment of DIV17 hippocampal dissociated neurons expressing mRuby3 (top panel) and PSD95.FingR-GFP (middle panel). PSD95 positive spines (blue arrow) and shaft PSDs (pink arrow) are indicated. 3D reconstruction (left top and middle panels) performed using the filament tracer tool in Imaris software. Identification of reconstructed PSD95 puncta was performed by using a diameter parameter of 0.15-0.8 µm with the spot detection tool. Dendritic body (gray), spines (light gray), and PSD95 puncta (red). (Bottom left panel) 90-degree forward rotation of 3D reconstruction of representative oblique section in image A. (Bottom right panel) 270-degree forward rotation of 3D reconstruction of representative oblique section in image A. **B)** Schematic experimental live-imaging setup used to induce long-term potentiation plasticity in hippocampal primary neurons.

